## Supplementary information for "The TUTase URT1 connects decapping activators and prevents the accumulation of excessively deadenylated mRNAs to avoid siRNA biogenesis"

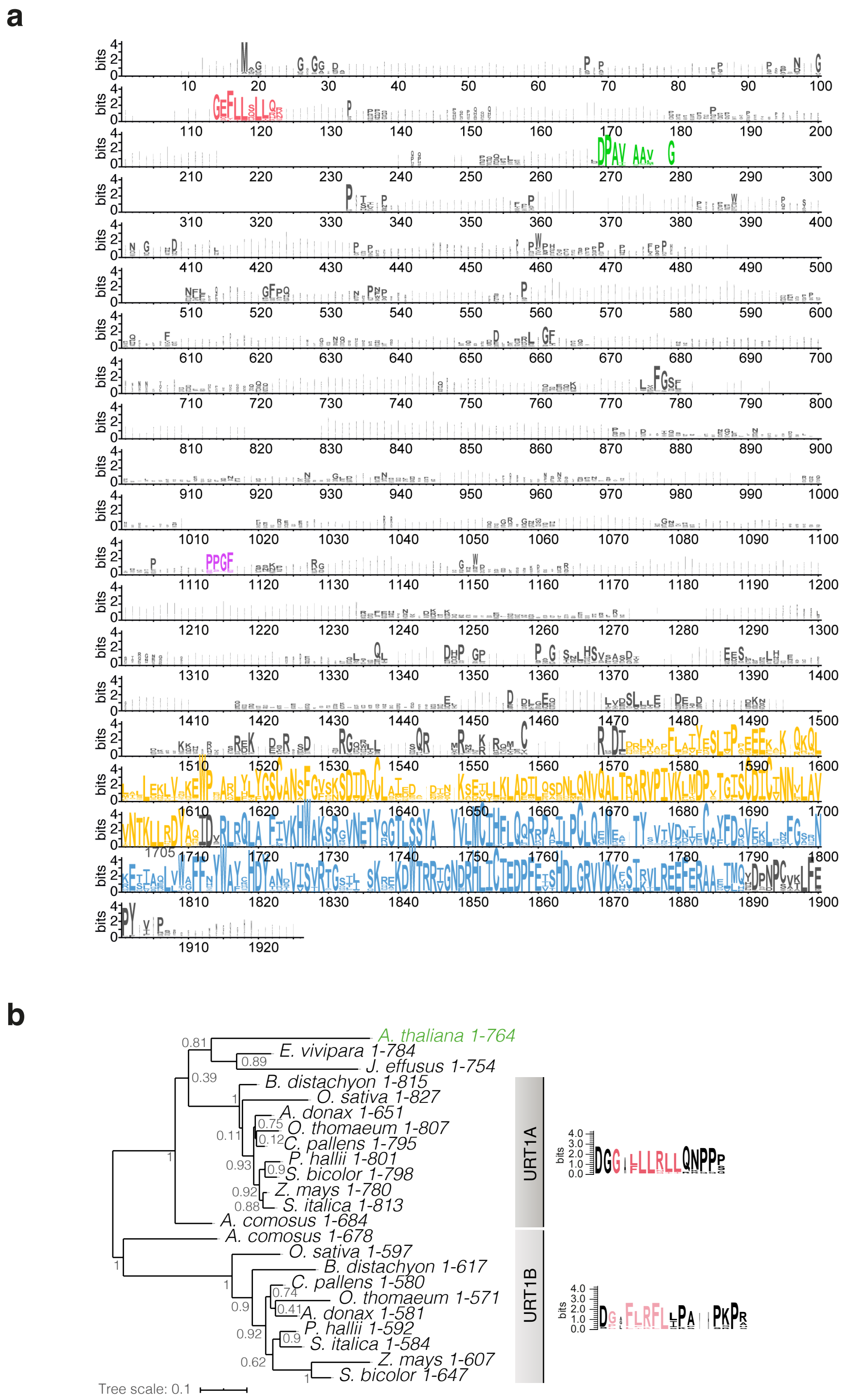


**Supplementary Fig. 1: Conservation of short linear motifs in URT1 sequences across land plants. a** Conservation of URT1 sequences across land plants. Consensus logo displayed in bits calculated from the alignment of 247 URT1 sequence homologs from land plants (see Supplementary Table 1). The conserved M1, M2 and PPGF motifs are highlighted in red, green and purple, respectively. β-like nucleotidyltransferase and poly(A) polymerase-associated domains are highlighted in yellow and blue, respectively. **b** Conservation of the M1 motif in Poales. URT1 homologous sequences in Poales and URT1 from *Arabidopsis thaliana* (see Supplementary Table 1) were aligned using Muscle, curated with Gblocks and used to construct a tree with the maximum-likelihood method and WAG substitution model implemented in PhyML (v. 3.1). Confidence values are shown on branches. The phylogenetic tree shows that URT1 homologs in Poales can be separated into 2 groups (A and B) with 2 distinct M1 motifs. The consensus logos showing the respective conservation of the M1 motif (red/light red in URT1A and URT1B groups, respectively) are shown on the right. The source data are available in Supplementary Table 1 and at [http://dx.doi.org/10.17632/ybcvvmtcn9.2].

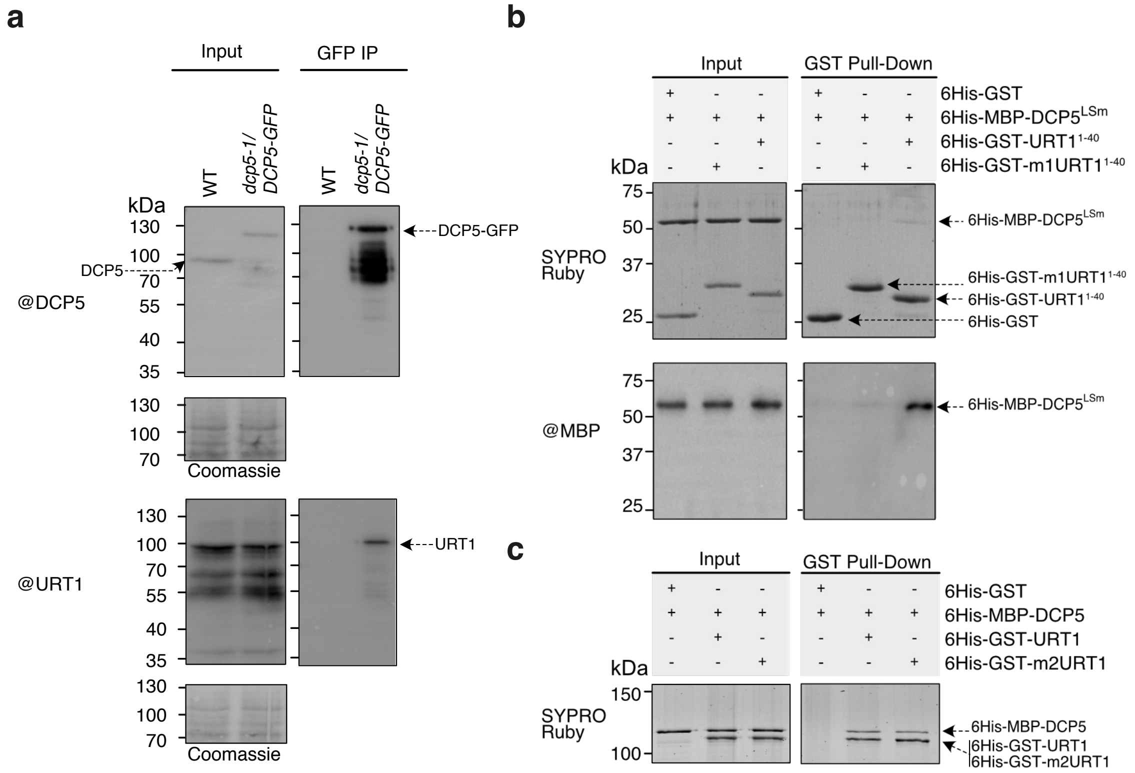


**Supplementary Fig. 2: Additional information related to the URT1-DCP5 interaction. a** URT1 co-purifies with DCP5-GFP expressed at endogenous level. WT plants and *dcp5-1* mutants expressing DCP5-GFP at endogenous level were used for IPs using anti-GFP antibodies. Input and eluates were probed with anti-DCP5 or anti-URT1 antibodies as indicated. **b** URT1's M1 motif and DCP5's LSm domain are sufficient for mediating the URT1-DCP5 interaction. *In vitro* GST pull-downs were performed in presence of RNase A using the indicated recombinant proteins. Pull-down and input fractions were analyzed by SDS-PAGE. Gels were either stained with SYPRO ruby or used for western analysis with anti-MBP antibodies to confirm the identity of 6His-MBP-DCP5$^{LSm}$. **c** URT1's M2 motif is not required for the URT1-DCP5 interaction. *In vitro* GST pull-downs were performed in presence of RNase A with the indicated recombinant proteins. Pull-down and input fractions were analyzed by SDS-PAGE and SYPRO ruby staining.

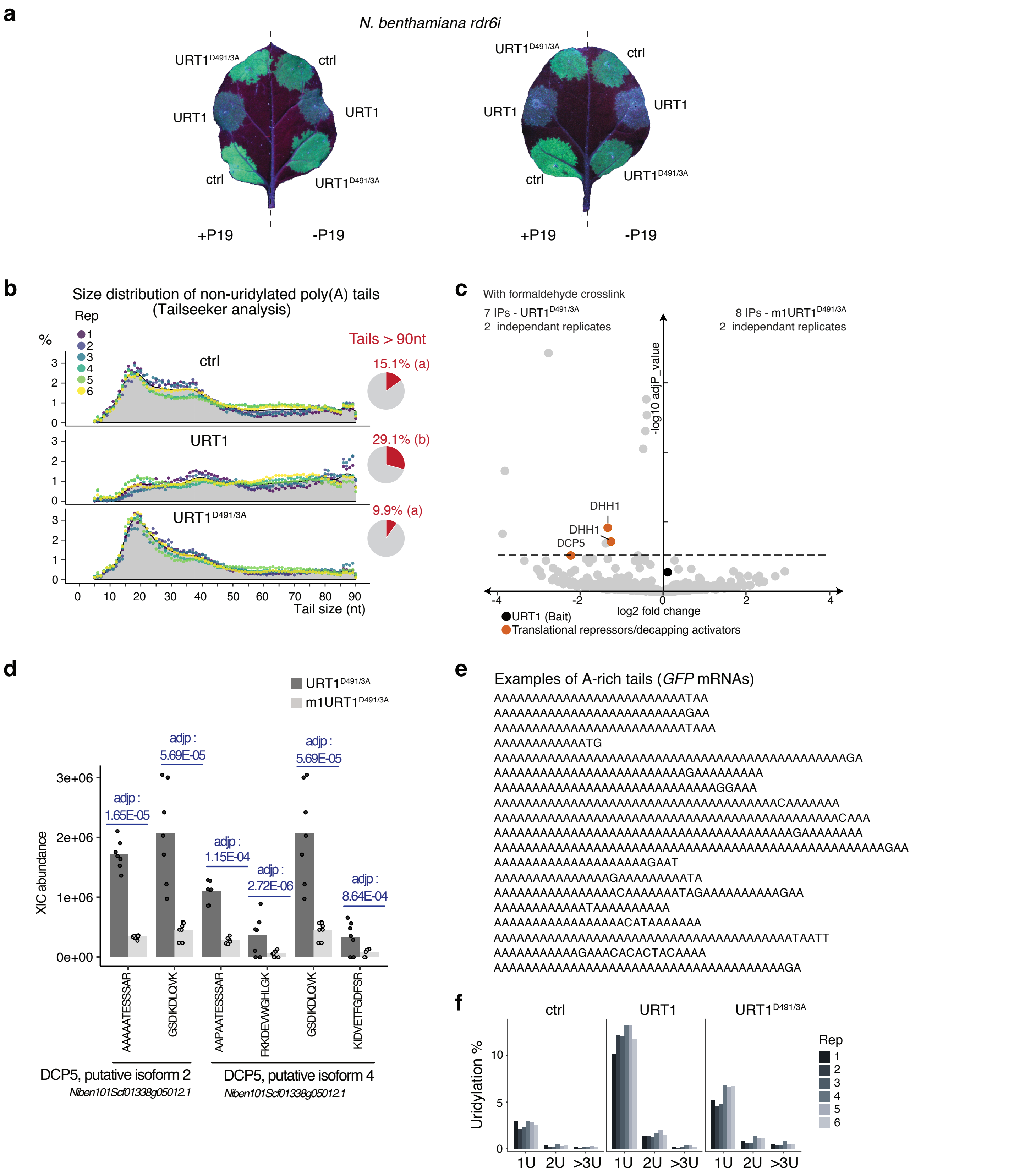


**Supplementary Fig. 3: Additional information related to URT1's impact on GFP expression. a** Picture of *N. benthamiana rdr6i* mutant plant leaves under UV light to detect the expression of the *GFP* reporter co-expressed with the different URT1-myc versions indicated, with or without co-expression of the P19 silencing suppressor. **b** Size distribution for non-uridylated poly(A) tails using the TAILseeker software (v3.1, https://github.com/hyeshik/tailseeker) for six biological replicates. Individual points are color-coded for each replicate and the average of all replicates is indicated as a grey area. The pie charts represent the average proportion of non-uridylated poly(A) tails longer than 90 As. Letters shown above pie charts represent significant statistical p-value (Wilcoxon rank-sum test, n=6). **c** Volcano plot showing *N. benthamiana* proteins differentially depleted (log2 fold change < 0) or enriched (log2 fold change > 0) in myc-tagged m1URT1$^{D491/3A}$ versus myc-tagged URT1$^{D491/3A}$ IPs. The dashed line indicates the significant threshold (adjusted p-value <0.05). **d** XIC (Extracted Ion Chromatograms)-based abundance for peptides that map *N. benthamiana* DCP5 isoforms. Bar plots show the average abundance of all replicates with individual points shown for each replicate. Adjusted p-values are indicated in blue for each peptide. XIC abundances and statistics (t-test) were determined using the Proteome Discover software. **e** Examples of A-rich tails for *GFP* mRNAs. **f** Percentage of poly(A) tails with 1U, 2U or at least 3U for six biological replicates. The source data are available in Supplementary Table 3 and at [http://dx.doi.org/10.17632/ybcvvmtcn9.2].

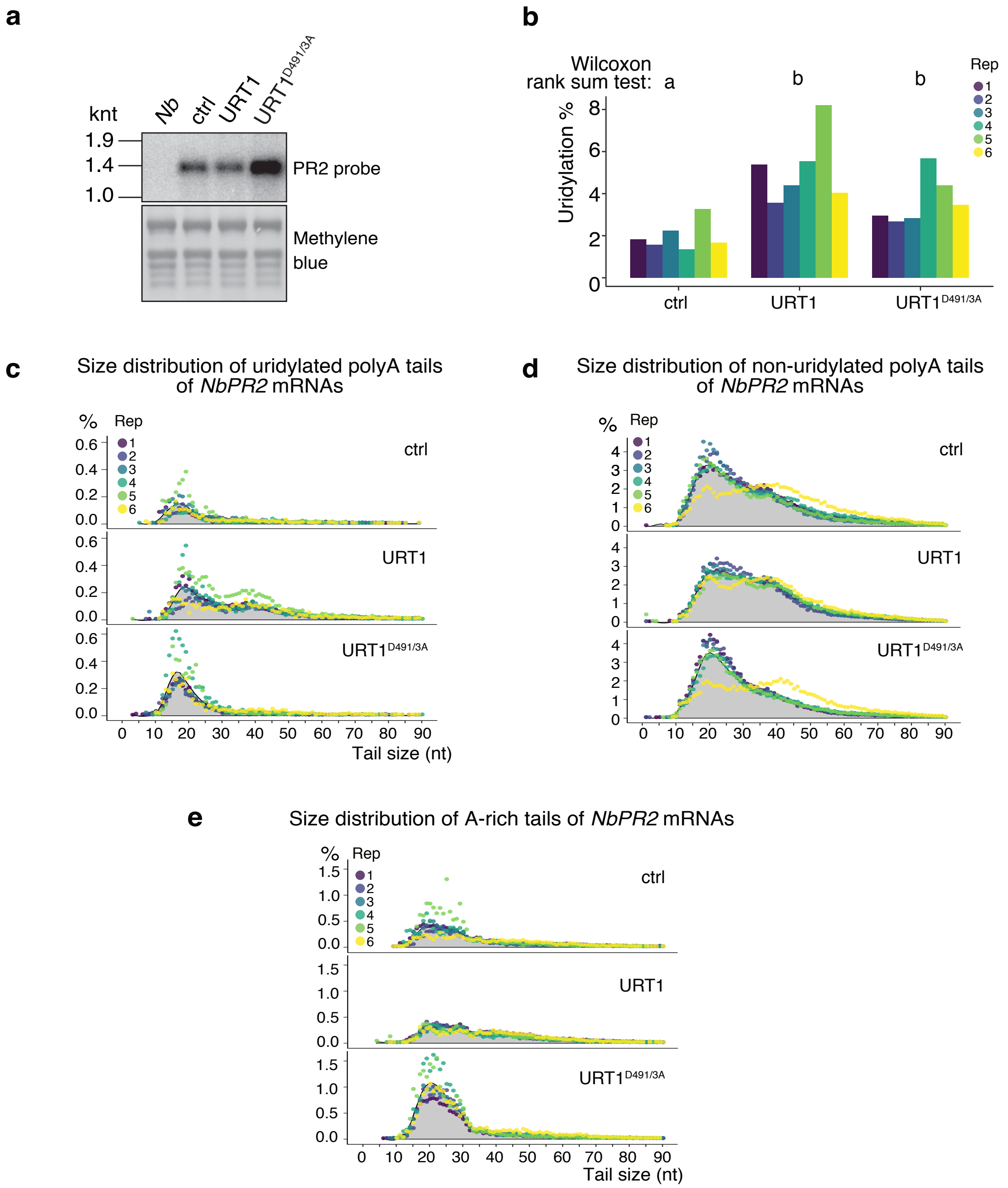


**Supplementary Fig. 4: URT1 ectopic expression affects poly(A) profiles of an endogenous mRNA.** **a** Northern blot showing the levels of endogenous *PR2* mRNAs upon ectopic URT1 expression. *Nb*: uninfiltrated *Nicotiana benthamiana* leaves. **b** Uridylation percentage of endogenous *PR2* mRNAs for six biological replicates. Letters represent significant statistical p-value (Wilcoxon rank-sum test, n=6). **c-e** Size distribution of poly(A) tails of endogenous *PR2* mRNAs. The percentages of sequences were calculated for six biological replicates for uridylated (**c**), non-uridylated (**d**) and A-rich (**e**) tails from 1 to 90 nucleotides. The percentages were calculated using the total number of sequences with tails from 1 to 90 nucleotides. Individual points are color-coded for each replicate and the average of all replicates is indicated as a grey area. The source data are available in Supplementary Table 3, at [https://www.ncbi.nlm.nih.gov/geo/query/acc.cgi?acc=GSE148409] and at [http://dx.doi.org/10.17632/ybcvvmtcn9.2].

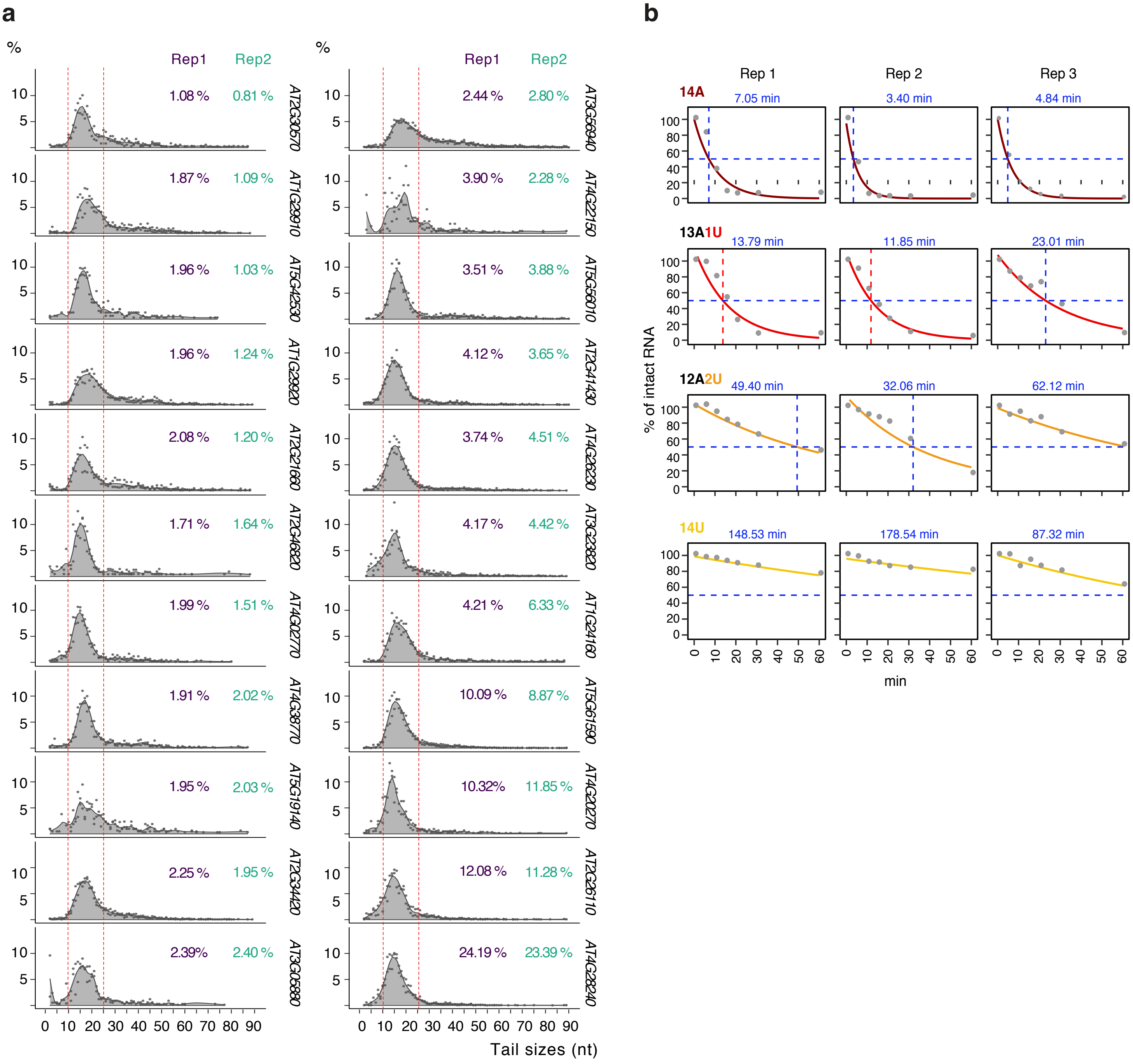


**Supplementary Fig. 5: Uridylation tags mostly oligoadenylated mRNAs with less than 25As and impedes deadenylation *in vitro*. a** Distribution of poly(A) tail sizes for uridylated sequences of 22 mRNAs in *Arabidopsis* plants. To facilitate intergenic profile comparison, the percentages of sequences were calculated using the total number of uridylated sequences for tails from 1 to 90 nt for two biological replicates. Tail length comprises As and 3' terminal uridines. Dashed lines indicate the 10 and 25 nt tail sizes. Numbers indicate the uridylation percentage in each WT replicate for each mRNA. **b** Half-life calculation for deadenylation assay. Points show the disappearance of the different RNA substrates after incubation with 6His-GST-CAF1b. Half-lives were calculated for each substrate and replicate using a quasi-Poisson regression (colored lines on graphs) and are indicated in blue. The source data are available in Supplementary Table 3, at [http://dx.doi.org/10.17632/ybcvvmtcn9.2], and at [https://www.ncbi.nlm.nih.gov/geo/query/acc.cgi?acc=GSE148406].

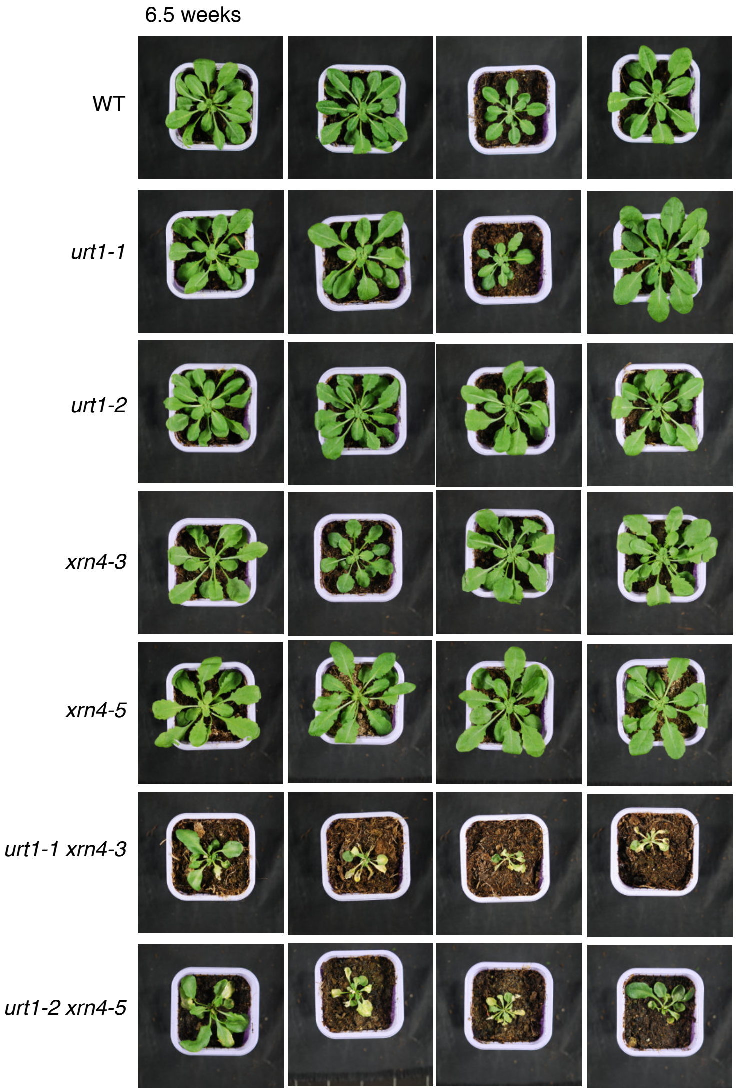


**Supplementary Fig. 6: Inactivation of both URT1 and XRN4 triggers a severe developmental phenotype.** Rosettes of WT, *urt1-1*, *urt1-2*, *xrn4-3*, *xrn4-5*, *urt1-1 xrn4-3*, *urt1-2 xrn4-5* plants. Plants were grown for 6.5 weeks in 12h light/12h darkness photoperiod conditions.

### Supplementary Methods

#### Characterization of URT1 sequence and phylogeny

The URT1 homologous sequences were collected by searching amino acid or nucleotide sequence databases using BLASTP or TBLASTN at National Center for Biotechnology Information (NCBI)[1] or Phytozome[2]. Nucleotide sequences were converted into protein sequences using the Expasy translate tool[3]. Positions of the different URT1 domains were determined using the Superfamily 2 webserver[4]. Amino acid compositional bias of the Arabidopsis URT1 sequence (Fig. 1a) was calculated using the fLPS program[5] with default parameter values (m = 15, M = 500, t = 0.001) using the amino-acid frequencies of Arabidopsis proteome (Araport 11) as a reference. Disorder scores shown in Fig. 1b and 1c were predicted using the ESpritz-NMR method[6] implemented in the FELLS webserver[7]. To compare URT1 homologs, protein lengths were normalized to calculate the disorder score per centile. Multiple protein alignments used in Fig. 1d, Supplementary Fig. 1a and 1b were constructed with the software MUSCLE (v3.8.31)[8] through the Jalview interface[9] using default parameters. Conservation scores of URT1 amino acids in Fig. 1d were calculated using the ConSurf webserver[10] with an alignment of 247 URT1 homolog sequences using Arabidopsis URT1 as a reference and the implemented Maximum Likelihood method[11] and WAG substitution model[12]. The sequence logo was generated using the Weblogo3 tool[13,14]. The phylogenetic tree of URT1 homologs in Poales (Supplementary Fig. 1b) was built with the maximum-likelihood method and WAG substitution model implemented in PhyML (v. 3.1) after curation of the alignment[15,16]. iTOL[17] was used to draw the three representation and calculate confidence values.

#### Bacterial expression of recombinant protein and purification

Recombinant proteins were expressed into *Escherichia coli* BL21 DE3 (genotype: F- ompT gal dcm lon hsdSB(rB- mB-) λ(DE3 [lacI lacUV5-T7 gene 1 ind1 sam7 nin5])) from constructs listed in Supplementary Table 7. Bacteria were grown in Luria-Bertani (LB) liquid medium at 37°C until an optical density at 600 nm (OD600) of 0.8 was reached. The temperature was then decreased to 20°C and protein expression was induced for 5 h by the addition of 1 mM of isopropyl-thiob-D-galactoside (IPTG). Bacteria were pelleted by centrifugation for 15 min at 4000 g, washed twice with 50 ml of PBS (137 mM NaCl, 10 mM $Na_2HPO_4$, 1.8mM $KH_2PO_4$, 2.7 mM KCl, pH 7.4). Bacteria were finally resuspended in lysis buffer (20 mM MOPS pH 7.2, 250 mM KCl, 15 % glycerol, 0.1 % Tween 20, 0.2 mM DTT and protease inhibitors (cOmplete, EDTA-free Protease Inhibitor Cocktail, Roche)) to obtain an OD600 of 20. Bacteria were disrupted by sonication cycles (2 s ON/5 s OFF for a total sonication time of 1 min) at an amplitude of 60 % (Vibracell Bioblock Scientific 75115). Cell debris were removed by centrifugation at 16,000 g for 30 min at 4°C. The bacterial lysate was filtrated through a 0.45 µm filter before incubation for 3 h with amylose resin (NEB) or glutathione-sepharose 4B resin (GE healthcare) at 4°C with constant rotation. The slurry was transferred to gravity-flow columns and washed with 20 mM MOPS pH 7.2, 250 mM KCl, 15 % glycerol and 0.1 % Tween 20. MBP tagged proteins were eluted from the amylose resin with an elution buffer composed of 20 mM MOPS pH 7.2, 150 mM KCl, 15 % glycerol, 0.1 % Tween 20 supplemented with 10 mM maltose monohydrate. For the purification of GST-tagged proteins, the elution buffer was supplemented with 10 mM reduced glutathione. The eluted protein fractions were dialysed overnight at 4°C using dialysis tubing cellulose membranes (Sigma-Aldrich, molecular weight cut-off of ~14,000 kDa) in 20 mM MOPS pH 7.2, 100 mM KCl, 15 % glycerol and 0.1 % Tween 20.

6His-MBP-DCP5 expression was performed in ZYP50502 autoinducing media[18]. Bacteria were grown at 37°C for 4 h and then at 20°C overnight. 650 ml of bacterial suspension were pelleted by centrifugation at 8,000 g for 30 min at 4°C and subsequently resuspended in a lysis buffer composed of Tris-HCl 50 mM pH 7.5, NaCl 300 mM, glycerol 5 % and lysed using a LM20 Microfluidizer® (Microfluidics). The cell extract was clarified by centrifugation at 17,000 g for 1 h at 4°C. The supernatant was filtered through a 0.22 µm filter before incubation for 1 h with amylose resin at 4°C with constant rotation. The resin-clarified lysate mix was applied on a gravity-flow column, and the resin was washed twice with 20 mL of lysis buffer. Bound proteins were eluted with 5 ml of Tris-HCl 50 mM pH 7.5, NaCl 300 mM, glycerol 10 % and 10 mM maltose. The elution fraction was dialyzed overnight at 4°C using dialysis tubing cellulose membranes (Sigma-Aldrich, molecular weight cut-off of ~14 000) in Tris-HCl 50 mM pH 7.5, NaCl 300 mM, glycerol 10 %. Full-length 6His-MBP-DCP5 was separated from a C-terminally truncated product by anion exchange chromatography using a MonoQ (Sigma-Aldrich) column and a linear gradient from 50 to 600 mM NaCl in 25 mM Tris-HCl pH7.5, 10 % glycerol.

**Quantification of GFP fluorescence in agroinfiltration experiments**

The intensity of the GFP fluorescence was quantified using ImageJ (v1.52). The images were first converted into 8-bit images and split into RGB single channels. For each leaf and infiltrated patch, the average pixel intensity of three different areas was measured on the green channel. To estimate the background level, average pixel intensity was also measured for three area outside infiltrated patches for each leaf. These background values were then subtracted from the average pixel intensity of each patch. Finally, the calculated fluorescence intensity was normalized to the intensity of the control patch of the corresponding leaf.

**Quantification of CAF1b activity**

Radiolabeled RNAs were visualized and quantified using an Amersham Typhoon IP Biomolecular Imager (GE Healthcare Life Sciences) and the QuanTL software. Three biological replicates were analyzed to quantify the disappearance of intact substrate RNA. Signals at time zero were used for normalization. Half-lives were calculated using the quasi-Poisson regression of the stats R packages (3.6.1).

**3’ RACE-seq data processing**

Fastq files were analyzed by a set of homemade scripts using python (v2.7), biopython (v1.63)[19] and regex (v2.4) libraries. Reads with low quality bases (= < Q10) within the 15-base random sequence of the read 2 or within the 30 bases downstream of the delimiter sequence were filtered out. Sequences with identical nucleotides in 15-base random sequence were deduplicated. Next, 20 nucleotides sequences corresponding to nucleotides of the transcript that maps downstream the forward PCR2 primer (Supplementary Table 6) were searched into reads 1 to identify the corresponding target mRNAs. One mismatch was tolerated. Matched reads 1 and their corresponding reads 2 were extracted and annotated. Reads 2 that contain the delimiter sequence were selected and subsequently trimmed from their random and delimiter sequences. Then, the analysis was divided into two steps. The aim of the first step was to identify the position of mRNA 3’ extremities and to detect untemplated nucleotides. To do this, the 30 nucleotide sequences downstream of the read 2 delimiter sequence were mapped to the corresponding reference sequence, which goes from the first nucleotide of the transcript that maps the forward PCR2 primer to the end of the mRNA. Up to four mismatches were tolerated, with the exception of the first five nucleotides downstream of the mapping site that had to perfectly map. To map the 3’ end position of reads 2 with untemplated tails, the sequences of the unmatched reads 2 were successively trimmed from their 3’ end, with a one nucleotide trimming step, until they could be mapped to the reference sequence or until a maximum of 30 nucleotide has been removed. For each successfully mapped read 2, untemplated nucleotides at the 3’ end were extracted. The aim of the second step was to analyze long mRNA poly(A) tail. Sequencing of long homopolymeric stretches causes a rapid decrease of sequencing quality, making it impossible to exactly map the 3’ end of mRNA with long poly(A). We thus looked for long T stretches of at least 10 Ts in the read 2 that failed to map the reference sequence. Poly(A) tails were searched with the constraint that it must begin in the first 30 cycles, which means that the maximal length of the added 3’ end modification is limited to 29 nucleotides. Finally, results from step 1 and 2 were compiled and the 3’ extension were analyzed. Python and bash source code are available as Mendeley data: http://dx.doi.org/10.17632/v8d9bd692c.1 (The link for reviewer access to the data is: https://data.mendeley.com/datasets/v8d9bd692c/draft?a=c426a729-7e3e-4099-bdf7-c08c87b1308d)

**Poly(A) length estimation using Tailseeker software**

We used the level 1 of the TAILseeker software (v3.1, https://github.com/hyeshik/tailseeker)[20] which performs PCR duplicate removal, poly(A) length measurement (≥5 nt) and identification of non-A additions to poly(A) tails. Default setting were used with exception of the following parameters “max_cctr_scan_left_space”, “max_cctr_scan_right_space” and “poly(A)_boundary_pos” that were set to 20, 20 and 120, respectively. The “third party basecaller” option was also disactivated. To identify the corresponding target mRNAs, 20 nt sequences that maps downstream the forward PCR 2 primer (Supplementary Table 6) were searched in the read 1. Finally, a homemade script based on python 2.7 was used to analyze the composition of non-A modifications detected by the Tailseeker software. Python and bash source code are available as Mendeley data: http://dx.doi.org/10.17632/v8d9bd692c.1

The link for reviewer access to the data is: https://data.mendeley.com/datasets/v8d9bd692c/draft?a=c426a729-7e3e-4099-bdf7-c08c87b1308d

**TAIL-seq data processing**

The base calls were retrieved after initial data processing by the MiSeq Control Software v 2.6 and the sequences having identical sequences in the 1st to 15th cycles in read 2 were deduplicated. In order to identify transcripts, read 1 sequences were mapped onto the *Arabidopsis thaliana* reference genome (TAIR 10) using Hisat2 (v2.1.0)[21] and Bowtie2[22]. The resulting alignment was annotated using the intersectBed tool from the BEDTools suite (v2.17.5)[23] and the Arabidopsis annotation file TAIR10_GFF3_genes.gff (http://www.arabidopsis.org). Reads 1 that map onto cytosolic mRNAs and their corresponding reads 2 were extracted and used for further analyses. Reads 2 that contain the delimiter sequence were selected and subsequently trimmed from their random and delimiter sequences. A homemade python script was then used to extract poly(A) tails (>5 nt) and their potential 3’ end modification from the remaining read 2. Poly(A) were detected with a constraint that it must begin within the first 30 cycles, so the maximum detectable 3’ end modification of poly(A) tails was limited to the last 29 nucleotides of insert. Detected poly(A) and 3’ extensions were finally analyzed for their size and composition using a homemade script based on python 2.7. Python and bash source code are available as Mendeley data: http://dx.doi.org/10.17632/v8d9bd692c.1

The link for reviewer access to the data is: https://data.mendeley.com/datasets/v8d9bd692c/draft?a=c426a729-7e3e-4099-bdf7-c08c87b1308d

**Small RNA-seq data processing**

Sequencing quality was assessed with a PhiX reference spiked in the flow-cell that gave an average sequencing error rate of 0.16 % with 92 % of the reference nucleotides having a Q-score >=30 (Phred calculation). ‘TGGAATTCTCGGGTGCCAAG’ sequence corresponding to the 20 nt sequence of the 5' end of the 3’ adapter was searched and removed using in-house Fasteris pipeline. Trimmed reads of 14 nt to 30nt in length were selected to be mapped to TAIR10 genome using Bowtie (v1.0.0)[24] by permitting up to 1 mismatch. Reads that could map to more than 50 loci were discarded and only the best alignment(s) of each small RNA sequence was kept. BEDTools suite (v2.17.0)[23] was then used to annotate and count reads that map to mRNAs using intersectBed and multiBamCov tools, respectively, with default settings. mRNAs were annotated based on TAIR10 annotation (http://www.arabidopsis.org). Differential analysis of small RNA accumulation between samples was performed using edgeR package (3.26.5) and its implemented negative binomial generalized log-linear model. P-value were adjusted using the Benjamini Hochberg method.
